# Aging reduces calreticulin expression and alters spontaneous calcium signals in astrocytic endfeet of the mouse dorsolateral striatum

**DOI:** 10.1101/2021.09.24.461710

**Authors:** Sara M. Zarate, Taylor E. Huntington, Pooneh Bagher, Rahul Srinivasan

**Author notes:** **Corresponding Author:** Rahul Srinivasan, Department of Neuroscience & Experimental Therapeutics, Texas A&M University College of Medicine, Bryan, TX 77807-3260, USA. **Author contributions:** SMZ performed all experiments and analyses, created figures, and contributed to writing and editing the manuscript. TEH created the tools and techniques to measure Ca^2+^ influx into mitochondria, and contributed to editing the manuscript. PB designed and supervised experiments and associated analysis and contributed to editing the manuscript. RS conceptualized, designed and coordinated the study, trained and supervised SMZ and TEH, designed and supervised experiments, advised on figures, provided resources and funding, and wrote the manuscript with SMZ. **CRediT author statement Sara M. Zarate:** Methodology, Validation, Formal analysis, Investigation, Writing – Original draft, Writing – Review & Editing, Visualization; **Taylor E. Huntington:** Methodology, Writing – Review & Editing; **Pooneh Bagher:** Conceptualization, Methodology, Formal Analysis, Writing – Review & Editing; **Rahul Srinivasan:** Conceptualization, Methodology, Validation, Formal Analysis, Investigation, Resources, Writing – Original draft, Writing – Review & Editing, Visualization, Supervision, Project administration, Funding acquisition.

## Abstract

Aging-related impairment of the blood brain barrier (BBB) and neurovascular unit (NVU) increases risk for neurodegeneration. Among the various cells participating in BBB and NVU function, spontaneous Ca^2+^ signals in astrocytic endfeet are crucial for maintaining BBB and NVU integrity. To assess if aging is associated with changes in spontaneous Ca^2+^ signals within astrocytic endfeet of the dorsolateral striatum (DLS), we expressed a genetically encoded Ca^2+^ indicator, Lck-GCaMP6f in DLS astrocytes of young (3-4 month) and aging (20-24 month) mice. Compared to young mice, endfeet in the DLS of aging mice demonstrated a decrease in calreticulin (CALR) expression, and dramatic alterations in the dynamics of endfoot membrane-associated and mitochondrial Ca^2+^ signals. While young mice required both extracellular and endoplasmic reticulum (ER) Ca^2+^ sources for generating endfoot Ca^2+^ signals, aging mice showed exclusive dependence on ER Ca^2+^. These data suggest that aging is associated with significant changes in Ca^2+^ buffers and Ca^2+^ signals within astrocytic endfeet, which has important implications for understanding mechanisms involved in aging-related impairment of the BBB and NVU.

**Graphical abstract:** 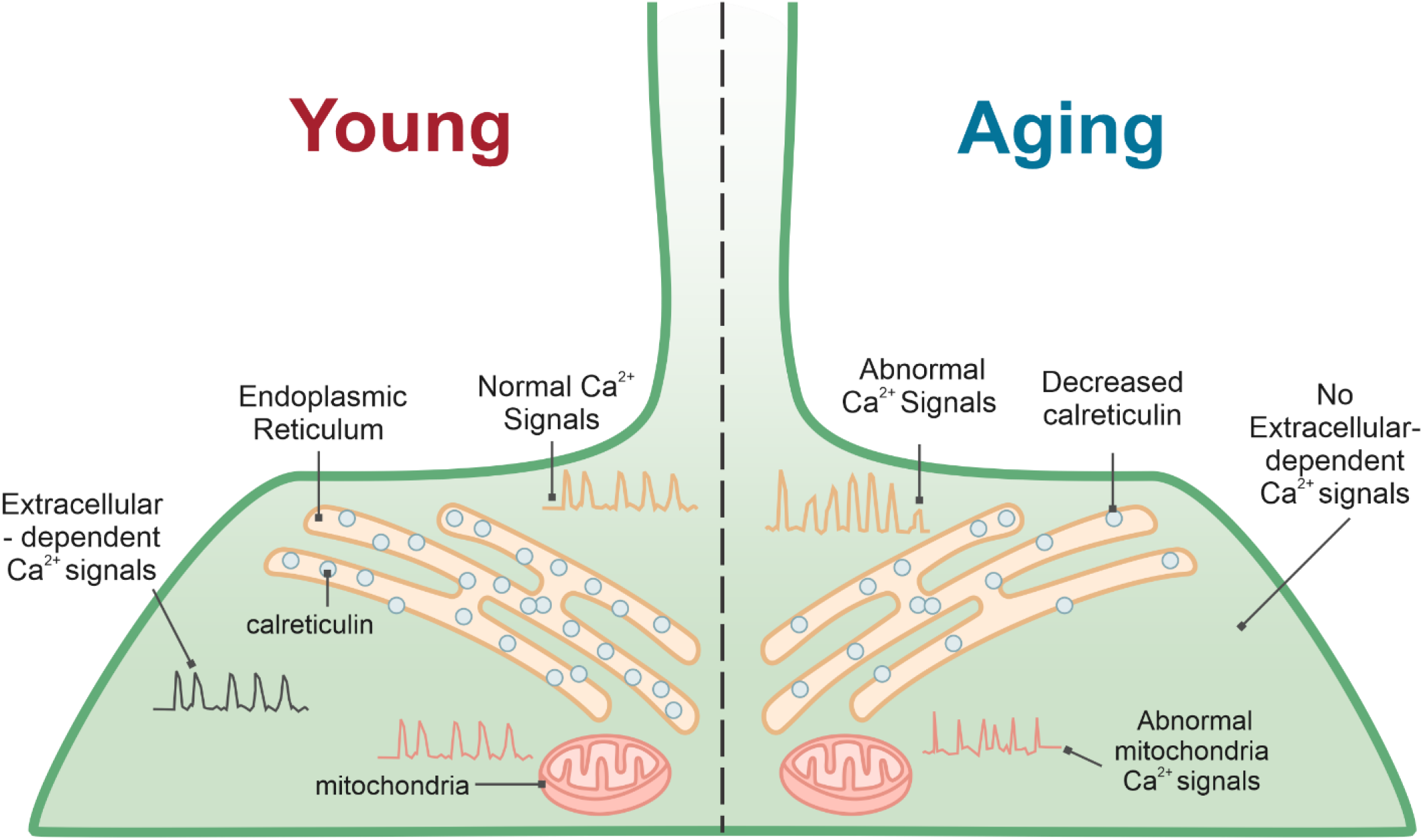

**Highlights:**

- Aging mice show reduced calreticulin expression in astrocytic endfeet
- Aging astrocytic endfeet show dramatic changes in spontaneous Ca^2+^ activity
- Ca^2+^ signals in aging endfeet depend exclusively on endoplasmic reticulum Ca^2+^

## Introduction

Advanced age is the single greatest risk factor for neurodegenerative conditions such as Parkinson’s disease (PD), Alzheimer’s disease (AD) (Hou et al., 2019), stroke (Leritz et al., 2011; Yousufuddin and Young, 2019), and dementia (Irwin et al., 2018). A large body of evidence strongly suggests that aging-related impairment of the blood brain barrier (BBB) and neurovascular unit (NVU) potentiates, and possibly triggers neurodegenerative processes in the brain (Cabezas et al., 2014; Cai et al., 2017; Govindpani et al., 2019; Grammas et al., 1999; Ott et al., 2018; Sweeney et al., 2019; Yu et al., 2020; Zhou et al., 2020). Among the multiple brain regions prone to neurodegeneration, the striatum is particularly vulnerable to neurovascular dysfunction and neurodegeneration because of its rich vasculature supply (Feekes and Cassell, 2006). This notion is supported by reports showing a higher percentage of lacunar infarcts in the striatum compared to other brain regions (Feekes et al., 2005), increased propensity for accumulating toxic striatal protein aggregates (Duda et al., 2002; Hanseeuw et al., 2019), and an increase in striatal BBB permeability during PD, AD and stroke (Gray and Woulfe, 2015; Haley and Lawrence, 2017; Sweeney et al., 2018). Furthermore, aging-related changes in the striatal BBB and NVU can lead to parkinsonian symptoms (Rosso et al., 2018), enlarged perivascular spaces (Chung et al., 2021), and PD due to dopaminergic neuron loss (Ivanidze et al., 2020). Although these studies indicate that striatal BBB and NVU dysfunction are important contributing factors for various forms of neurodegeneration, we know very little about the specific molecular mechanisms involved in altering BBB and NVU function during aging.

The BBB and NVU are comprised of distinct cellular elements, *viz*., endothelial cells, pericytes and astrocytes, each of which plays important and specific roles in neurovascular function (Garcia and Longden, 2020; Mishra et al., 2016; Pandit et al., 2020; Villabona-Rueda et al., 2019). Among these cell types, astrocytes are unique in that they simultaneously contact neurons and the vasculature, thus enabling these cells to regulate blood flow in response to brain activity (Attwell et al., 2010; Belanger et al., 2011; Heithoff et al., 2021; Howarth, 2014). Astrocytes perform this vital function via spontaneous Ca^2+^ signals within endfeet that completely ensheath capillaries (Girouard et al., 2010; Zhang et al., 2019). Indeed, Ca^2+^ signals in astrocytic endfeet have been shown to mediate vascular repair after injury (Gobel et al., 2020), regulate brain volume (Eilert-Olsen et al., 2019), as well as maintain NVU coupling (Dunn et al., 2013). Thus, at a functional level, astrocytic endfoot Ca^2+^ signals govern critical aspects of BBB integrity and NVU function. It follows that aging-related changes in striatal astrocytic endfoot Ca^2+^ signals likely contribute to BBB and NVU dysfunction during neurodegeneration. Based on the strong body of evidence pointing to striatal vascular dysfunction in various forms of aging-related neurodegeneration as well as a central role for astrocytic endfoot Ca^2+^ signals in maintaining BBB integrity and NVU function, we asked if aging is associated with alterations in spontaneous astrocytic endfoot Ca^2+^ events within the dorsolateral striatum (DLS) of acutely extracted live brain slices from young and aging mice.

We show that when compared to young mice, DLS brain slices of aging mice display dramatic alterations in the kinetics, dynamics, and sources of spontaneous membrane-associated Ca^2+^ events, as well as changes in Ca^2+^ influx into astrocytic endfoot mitochondria. These aging-related alterations in endfoot Ca^2+^ events are associated with significant reductions in the expression of a major endoplasmic reticulum (ER) Ca^2+^ buffering protein, calreticulin (CALR). Our findings have important implications for understanding how an aging-related reduction in CALR expression can alter astrocytic endfoot Ca^2+^ signals in the BBB and NVU, eventually resulting in vascular dysfunction during aging and aging-related neurodegeneration.

## Materials and Methods

### Mice

Male and female C57BL/6J mice (Jackson laboratory, Bar Harbor, ME, USA) were used for all experiments. Young mice were 3-4 months of age, and aging mice were 20-24 months of age. Mice were aged in house, in the animal vivarium at Texas A&M University. All experiments were performed in accordance with Texas A&M University IACUC regulations and protocols. Food and water were provided *ad libitum*. All mice were maintained on a 12 h light-dark cycle.

### Adeno-associated virus (AAV) injection into the DLS of mice

Young and aging mice were deeply anesthetized using isoflurane dispensed from a SomnoSuite Low Flow Anesthesia System (Kent Scientific, Torrington, CT), and a craniotomy was performed as previously described (Huntington and Srinivasan, 2021; Jiang et al., 2014). AAVs were injected into the right DLS using a custom-made glass injection pipette with a 6 μm tip diameter, at a rate of 750 nl/min using a Harvard Apparatus Pump 11 Pico Plus Elite, 70-41506 (Harvard Apparatus, Holliston, MA). To image membrane-associated astrocyte endfoot Ca^2+^ events in young and aging mice, 1 μl of AAV2/5-GfaABC1D-Lck-GCaMP6f (10^13^ genome copies/ml) (Addgene viral prep # 52924-AAV5) was injected into the DLS. The Lck motif at the N-terminal end of GCaMP6f enables plasma-membrane insertion of GCaMP, and visualization of membrane-associate Ca^2+^ signals (Srinivasan et al., 2016). To image astrocyte endfoot mitochondrial Ca^2+^ events in young and aging mice, 2 μl of AAV2/5-GfaABC1D-mito7-GCaMP6f (10^13^ genome copies/ml) (Huntington and Srinivasan, 2021) was injected into the DLS. Coordinates for stereotaxic injection of AAVs were 0.8 mm rostral to the bregma, 2 mm lateral to the midline and 2.4 mm ventral to the pial surface.

### Confocal imaging of acute brain slices

Young and aging mice were deeply anesthetized with isoflurane and the left ventricle of the heart was rapidly accessed by opening the chest cavity. 200 μl of DyLight 594 labeled *Lycopersicon esculentum* lectin (tomato lectin, TL) (DL-1177, Vector Laboratories, Burlingame, CA) was injected into the apex of the beating left ventricle. 2 – 3 min following TL injection into the left ventricle, the mouse was decapitated and the brain was extracted, then blocked within 2 min. for obtaining live brain slices. 400 µm coronal striatal slices were cut using a Microslicer 01N (Ted Pella) in a solution comprising 194 mM sucrose, 30 mM NaCl, 4.5 mM KCl, 1.2 mM NaH_2_PO_4_, 26 mM NaHCO_3_, 10 mM D-glucose, and 1 mM MgCl_2_ bubbled with 95% O_2_ and 5% CO_2_. Live brain slices were incubated in artificial cerebrospinal fluid (aCSF) composed of 126 mM NaCl, 2.5 mM KCl, 1.24 mM NaH_2_PO_4_, 1.3 mM MgCl_2_, 2.4 mM CaCl_2_, 26 mM NaHCO_3_, 10 mM D-glucose at 33°C for 30 min and then maintained at 23°C in aCSF for the duration of the experiment.

Live mouse striatal brain slices were imaged with an Olympus FV1200 upright laser-scanning confocal microscope using a 40x water immersion objective lens (NA 0.8), and a digital zoom of 3x. We used a 488 nm excitation wavelength at 10% of maximum intensity of a 100 mW Argon laser to record membrane-associated astrocyte endfoot and mitochondrial Ca^2+^ events. A 25 mW HeNe 594 nm excitation laser line at 10% of maximum intensity was used to visualize blood vessels (BVs) and localize astrocytic endfoot Ca^2+^ events in the DLS. All BVs reported in this study were capillaries with a diameter range of 5 to 10 μm. Endfoot membrane-associated and mitochondrial Ca^2+^ events in close proximity to TL-labeled BVs were identified as astrocytic endfoot Ca^2+^ events and imaged for 10 min at 1 frame per second (FPS). Imaging parameters (laser intensity, HV, gain, offset, aperture diameter) were maintained constant across all imaging sessions.

Imaging of Ca^2+^ events in astrocytic endfeet was performed by first imaging spontaneous Ca^2+^ events near BVs, followed by sequential depletion of extracellular Ca^2+^ and ER Ca^2+^ in the slice. To deplete extracellular Ca^2+^, spontaneous Ca^2+^ events were recorded for 5 min in bath perfused aCSF, followed by bath perfusion of zero Ca^2+^ aCSF, in which CaCl_2_ was omitted. To deplete ER Ca^2+^ stores, 20 µM cyclopiazonic acid (CPA, Abcam, Cambridge, MA, ab120300) was perfused into the bath for 15 min, after which Ca^2+^ events were recorded for 5 min from the same field of view.

### Immunohistochemistry

Mice were deeply anesthetized with isoflurane and transcardially perfused with 1X PBS, followed by 10% formalin. Brains were extracted and stored in 10% formalin for 48 h at 4°C then dehydrated in 30% sucrose in 1X PBS (Sigma, St Louis, MO, cat# S7903) for 48 h. A Microm HM 550 cryostat was used to cut 40 µm coronal sections of the striatum that were stored at 4°C in 0.01% sodium azide (Sigma, cat# S2002) until the day of immunostaining. Sections were washed 3x in 1X PBS, then permeabilized and blocked in 0.5% Triton X-100 and 10% normal goat serum at room temperature for 1 h. To prevent cross reactivity, sections were sequentially stained with CALR and AQP4 primary and secondary antibodies. Sections were first incubated overnight at 4°C in rabbit anti-CALR (1:500, ThermoFisher, Waltham, MA, cat# PA3900) then goat anti-rabbit Alexa Fluor 488 (1:2000, Abcam, cat# ab150077) secondary antibody for 2 h at room temperature. After 3 washes in 1X PBS, sections were stained with rabbit anti-AQP4 (1:200, Alomone, cat# AQP-004) overnight at 4°C, followed by goat anti-rabbit Alexa Fluor 594 (1:2000, Abcam, cat# ab150176) for 2 h at room temperature. Following staining, sections were mounted on glass slides and imaged using an FV1200 inverted confocal microscope equipped with a 60x oil immersion objective.

### Analysis of Ca^2+^ events in astrocytic endfeet

Membrane-associated and mitochondrial astrocytic endfoot Ca^2+^ events were identified as Ca^2+^ events occurring immediately adjacent to the abluminal side of TL-labeled DLS BVs. Regions of interest (ROIs) for endfoot Ca^2+^ events identified in this way were generated as previously described, using the ImageJ plugin, GECIquant (Srinivasan et al., 2015). dF/F traces of GCaMP6f fluorescent signals were generated from the acquired ROIs and analyzed with Minianlysis 6.0.07 (Synaptosoft) to generate frequency (events/min), amplitude (dF/F), and half-width (s) values for each trace.

For expanding endfoot Ca^2+^ waves, the velocity (µm/s), distance traveled (µm), and area (µm^2^) of membrane-associated astrocytic endfoot Ca^2+^ events were analyzed using ImageJ. All confocal movies for analysis of these parameters were acquired at a frame rate of 1 FPS. For analyzing velocity, the distance covered by endfoot Ca^2+^ events was determined by using the line tool in ImageJ to trace the length of Ca^2+^ events along the corresponding BV across multiple frames, then divided by time. Distance traveled corresponds to the maximum length attained by any given Ca^2+^ event along a BV. The area of propagating Ca^2+^ events was quantified using the polygon tool in ImageJ. Using the polygon tool, the maximum area attained by a propagating Ca^2+^ event along the BV was manually outlined, and the area was measured.

### Sampling and statistics

Individual groups consist of all the DLS endfoot ROIs from 3 to 7 mice. Statistics were performed using Origin 2019 (OriginLab, Northampton, MA). For statistical testing, datasets were first tested for normality. Non-normal datasets were subjected to either a Mann-Whitney or paired sample Wilcoxon signed rank test. A two-sample t test was used to compare normally distributed datasets. *p* < 0.05 was considered statistically significant. Sample sizes, statistical tests used, and exact p-values are reported in figures and figure legends.

## Results

### Aging alters multiple parameters of spontaneous membrane-associated astrocytic endfoot Ca^2+^ events in the DLS

To measure spontaneous membrane-associated astrocytic endfoot Ca^2+^ events, the DLS of either young or aging mice was injected with an AAV expressing membrane localized GCaMP6f (Lck-GCaMP6f), driven by an astrocyte-specific GfaABC1D promoter (AAV2/5-GfaABC1D-Lck-GCaMP6f). Intraventricular administration of red fluorescent TL just prior to live brain slicing enabled visualization of BVs in the DLS (Figure 1A), while Lck-GCaMP6f was used to visualize astrocytic endfoot Ca^2+^ events (Figure 1B). Confocal imaging of live striatal brain slices co-labeled with TL in BVs and Lck-GCaMP6f in astrocytic endfeet revealed that both young and aging mice exhibited robust spontaneous endfoot Ca^2+^ events in ROIs immediately adjacent to TL labeled BVs in the DLS (Figures 1B and 1C; supplementary movies 1 and 2).

**Figure 1.**
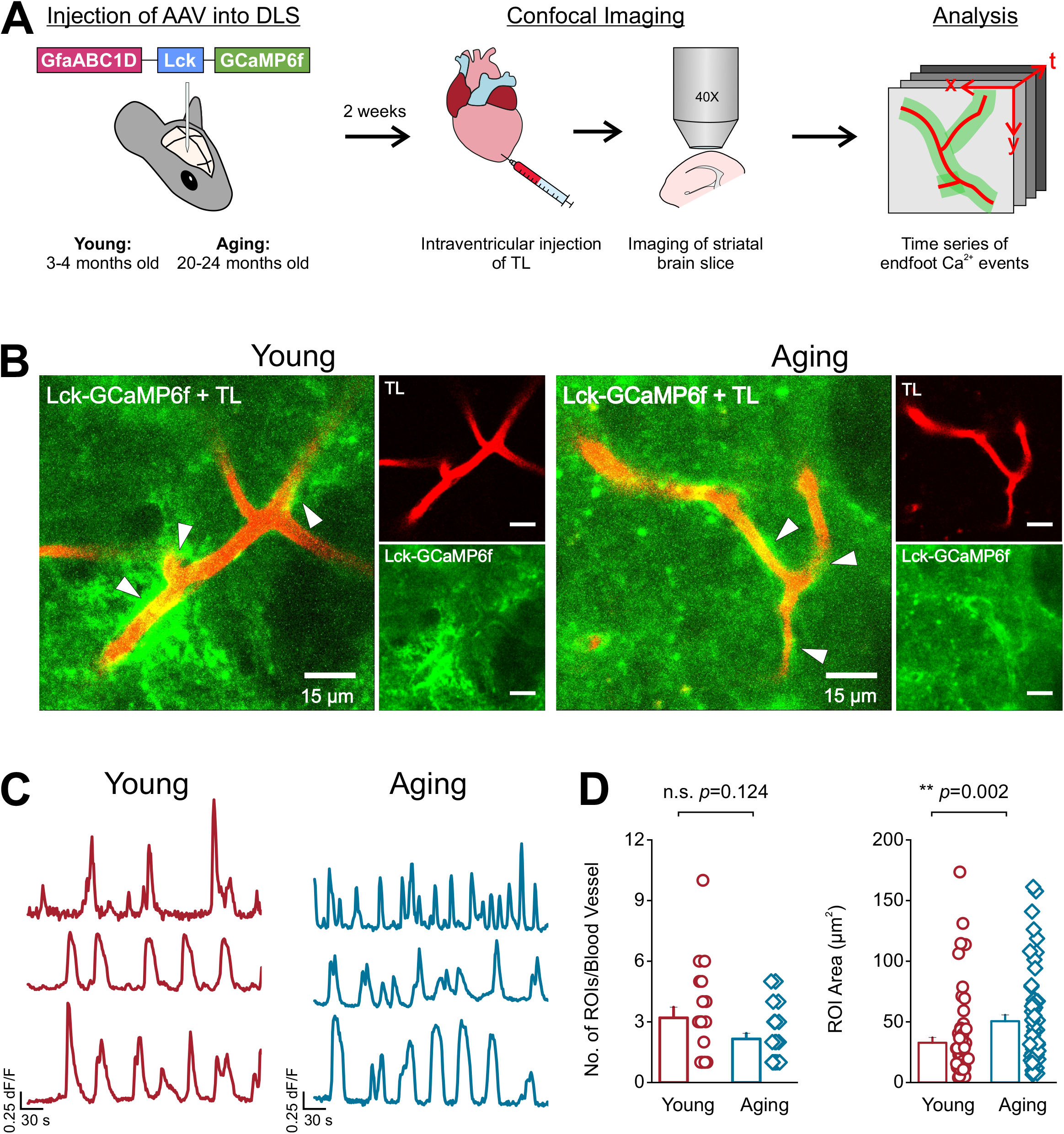
Astrocyte endfoot expression of Lck-GCaMP6f in young and aging mice. (**A**) AAV-GfaABC1D-Lck-GCaMP6f was injected into the DLS of young and aging mice. 2 weeks later, intraventricular injection of tomato lectin (TL) was performed, and striatal brain sections were collected for recording and measuring endfoot Ca^2+^ events in the DLS. (**B**) Representative images of young and aging astrocyte endfeet expressing Lck-GCaMP6f (green) immediately adjacent to TL labeled blood vessels (red) in the DLS, scale bar = 15 μm. White arrows point to areas where endfoot Ca^2+^ events initiated. (**C**) Example traces of endfoot Ca^2+^ events recorded from young and aging mice. (**D**) Comparisons of the number (left) and area (right) of ROIs generated in young and aging mice. For young mice, n = 64 ROIs and 20 blood vessels from 5 mice. For aging mice, n = 57 ROIs and 26 blood vessels from 7 mice. Error bars are S.E.M and all *p* values are based on Mann-Whitney test.

We found that the number of endfoot Ca^2+^ ROIs was greater in young mice when compared to aging mice (3.2 ± 0.53 ROIs per BV in young and 2.15 ± 0.28 ROIs per BV in aging mice), while the average area of endfoot Ca^2+^ ROIs in aging mice was ∼36% larger than in young mice (ROI area = 32.79 ± 4.3 µm^2^ in young and 50.6 ± 5.15 µm^2^ in aging mice) (Figure 1D). Next, we measured the kinetics (frequency, amplitude, and half width) of DLS astrocytic endfoot Ca^2+^ events in young and aging mice. Young mice demonstrated an average endfoot Ca^2+^ event frequency of 1 event/min, which was 20% faster than aging mice (Figure 2A). Cumulative frequency distributions of endfoot Ca^2+^ events in young mice revealed that 90% of events occurred between 0.2 and 1.6 events/min that segregated into 8 regularly spaced intervals. By contrast, in aging mice, 80% of all Ca^2+^ events occurred between 0.2 and 1.2 events/min, and segregated into more than 20 smaller spaced intervals (Figure 2B). Endfoot Ca^2+^ events in young mice displayed a constant frequency interval of 0.2 events/min, while endfoot Ca^2+^ events in aging mice showed a significantly shorter interval of ∼0.1 events/min, which was highly variable (Figure 2C). Aging mice also showed a 67% greater increase in Ca^2+^ event amplitude and a 40% greater increase in Ca^2+^ event half width when compared to young mice (Figure 2D).

**Figure 2.**
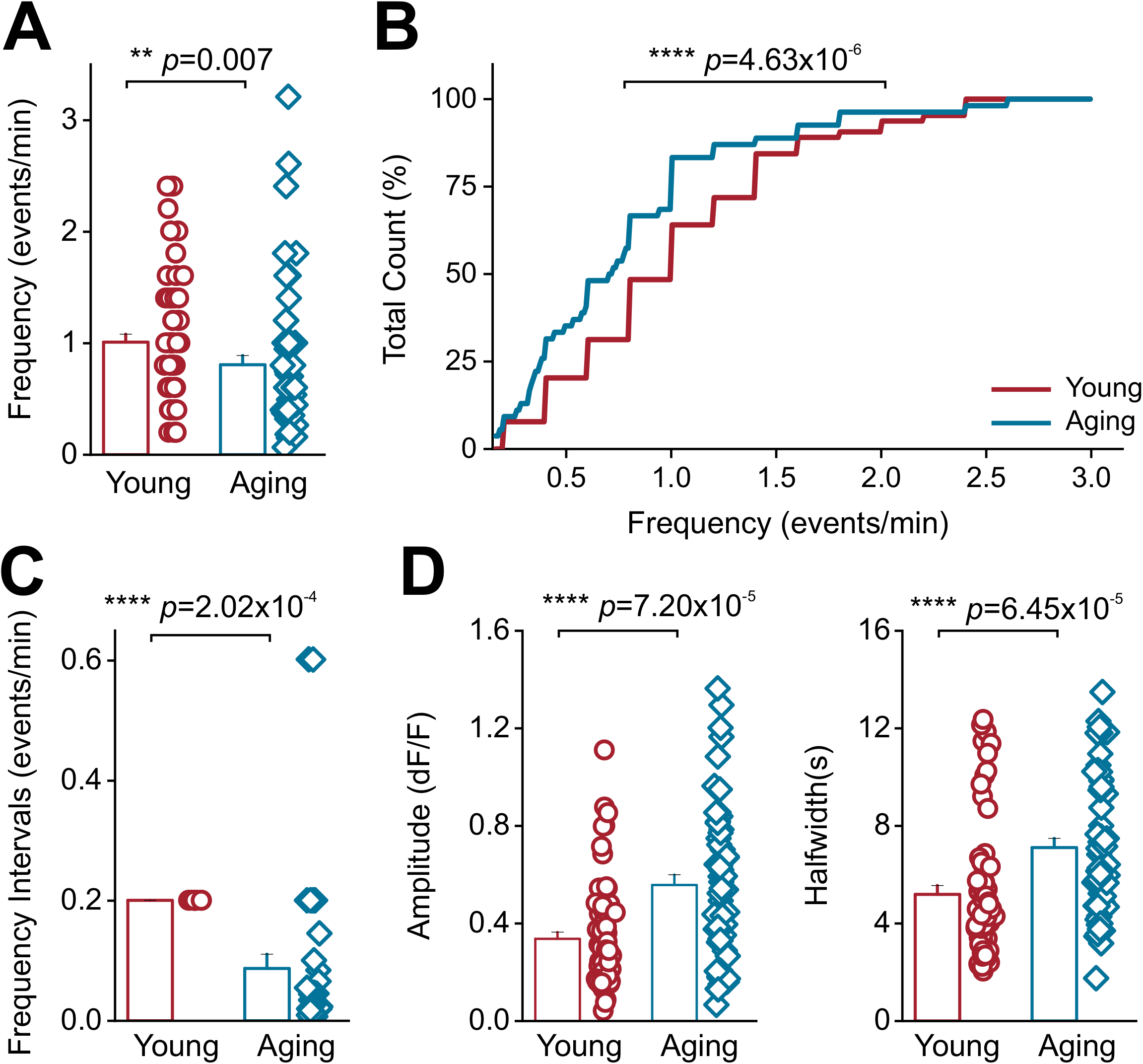
Aging alters Ca^2+^ event kinetics. (**A**) Plot of Ca^2+^ event frequencies in the endfeet of young and aging mice (**B**) Cumulative distribution of frequencies in young and aging mice (**C**) Plot of frequency intervals in young and aging mice (**D**) Amplitude (left) and half width (right) of spontaneous Ca^2+^ events in young and aging mice. For young mice, n = 64 ROIs and 20 blood vessels from 5 mice. For aged mice, n = 57 ROIs and 26 blood vessels from 7 mice. Error bars are S.E.M and for panels A, C, and D *p* values are based on Mann-Whitney tests. For the cumulative frequency plot in panel B, the *p* value is based on a Kolmogorov-Smirnov test.

Endfoot Ca^2+^ events in young and aging mice occurred as expanding waves traveling along the length of BVs (Figure 3A; supplementary movies 1 and 2). For these expanding Ca^2+^ waves, the velocity, distance, and area covered were measured. Compared to young mice, aging mice showed a significant increase in the expansion velocity of endfoot Ca^2+^ events (10.98 ± 1.45 µm/s in aging mice and 5.77 ± 0.93 µm/s in young mice) but showed no differences either in the distance covered by Ca^2+^ waves or the total area of Ca^2+^ waves (Figure 3B).

**Figure 3.**
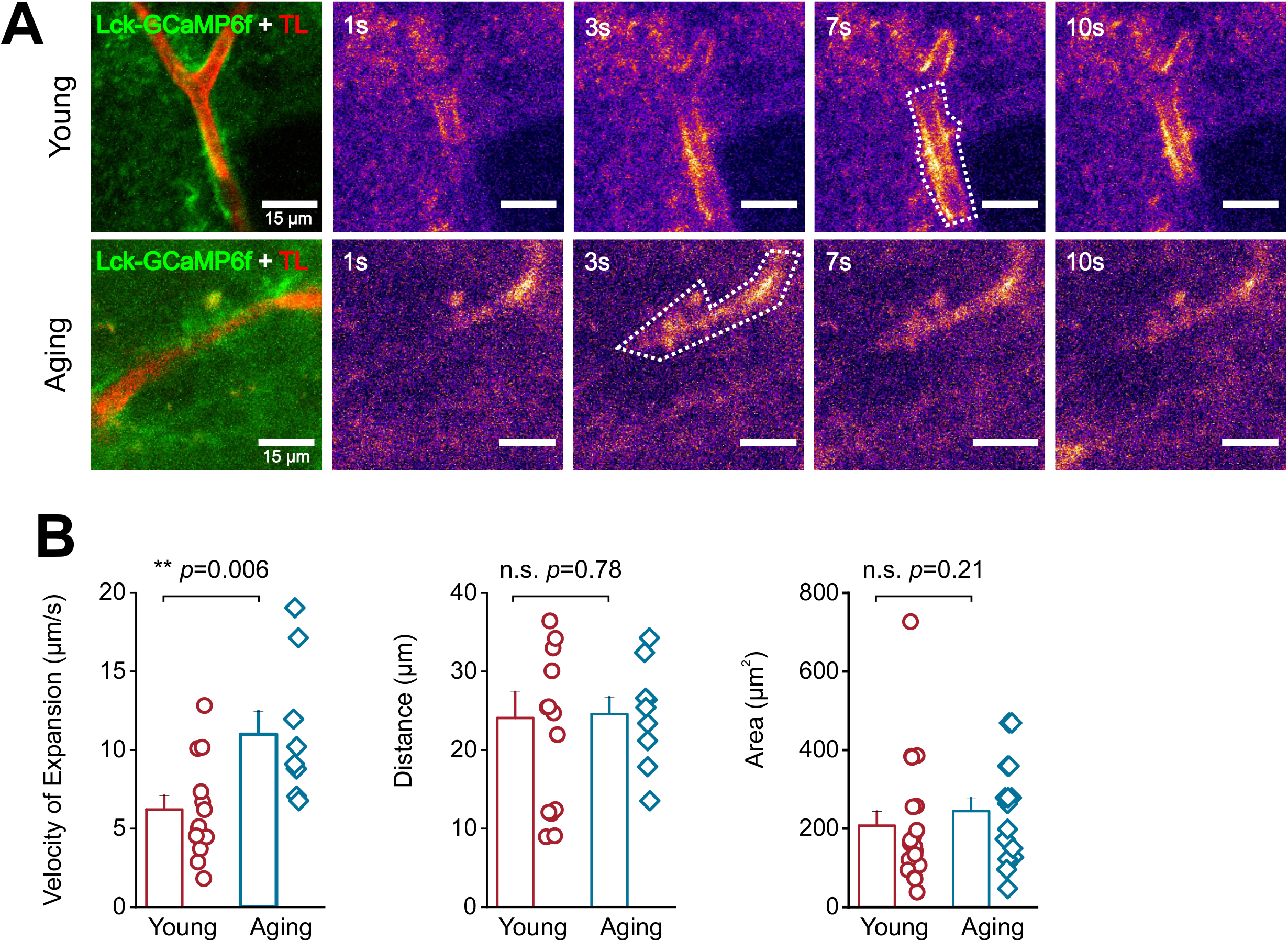
Aging increases the velocity of expanding Ca^2+^ waves in astrocyte endfeet. (**A**) A representative t-stack of an expanding endfoot Ca^2+^ wave in the DLS of a young (top) and aging (bottom) mouse is shown. Panels to the right of t-stacks show representative pseudo-colored time lapse images of endfoot Ca^2+^ waves for young (top) and aging (bottom) mice. The white dotted lines in time-lapse images outline the maximum area attained by expanding endfoot Ca^2+^ waves, scale bar = 15 μm (**B**) Bar graphs showing the velocity (left), distance traveled (middle), and area (right) of endfoot Ca^2+^ waves. For young mice, n = 14 ROIs and 6 blood vessels from 3 mice. For aging mice, n = 9 ROIs and 4 blood vessels from 3 mice Error bars are S.E.M and all *p* values are based on Mann-Whitney tests.

Taken together, these data show that aging significantly alters multiple aspects of the kinetics of spontaneous endfoot Ca^2+^ events in the DLS.

### Aging increases the dependence of astrocytic endfoot signals on ER Ca^2+^ stores

Having found that the DLS of aging mice show significant alterations in spontaneous astrocytic endfoot Ca^2+^ signals, we sought to determine if there were also differences in the extent to which endfoot Ca^2+^ events in young versus aging mice depend on extracellular and intracellular Ca^2+^ stores.

Extracellular and intracellular Ca^2+^ stores were sequentially depleted and astrocytic endfoot Ca^2+^ events were recorded. In young mice, depletion of extracellular Ca^2+^ with zero Ca^2+^ aCSF caused a significant 23% decrease in Ca^2+^ event frequency (Figure 4A and B; supplemental movie 1) with no effect on the amplitude and half width of Ca^2+^ events (Figure 5A). Depletion of ER Ca^2+^ stores in young mouse brain slices with 20 µM of the Ca^2+^-ATPase SERCA pump inhibitor, CPA, for 15 minutes dramatically reduced Ca^2+^ event frequency by ∼40% (Figure 4A and B). In addition, young mice showed a 78% decrease in the amplitude and a 43% decrease in the half width of Ca^2+^ events that persisted after CPA exposure (Figure 5C).

**Figure 4.**
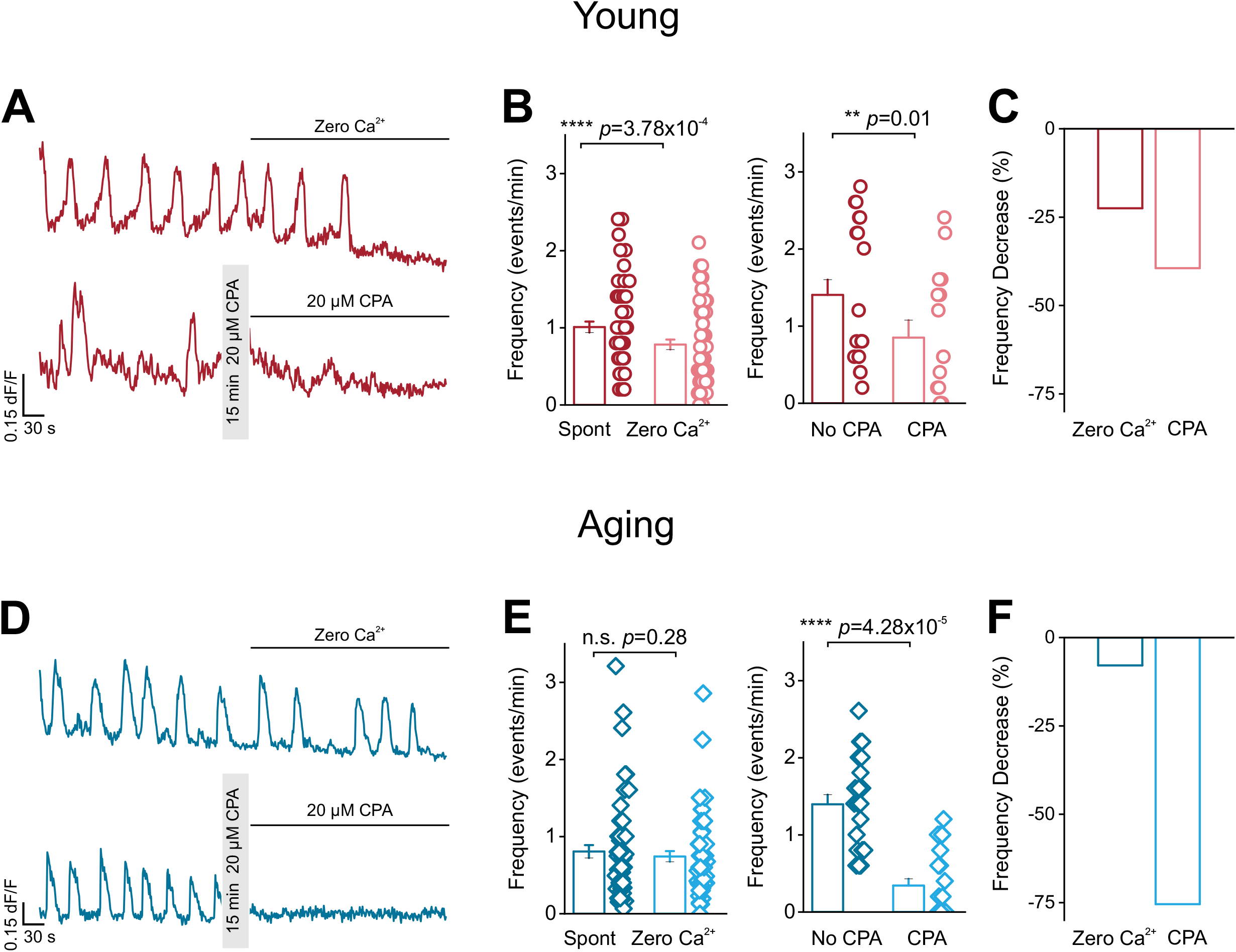
Astrocyte endfeet in aging mice exclusively rely on ER Ca^2+^ stores to generate spontaneous Ca^2+^ events. (**A**) Representative endfoot Ca^2+^ event traces from the DLS of young mice after bath application of zero Ca^2+^ aCSF or after 15 min incubation with 20 µM CPA (**B**) Bar graphs showing the average Ca^2+^ event frequency for young mice after bath application of zero Ca^2+^ aCSF (left graph) or after 15 min incubation with 20 µM CPA (right graph) (**C**) Bar graph of average % decrease in Ca^2+^ event frequency in young mice exposed to either zero Ca^2+^ aCSF or CPA (**D**) Representative endfoot Ca^2+^ event traces from the DLS of aging mice after bath application of zero Ca^2+^ aCSF or after 15 min incubation with 20 µM CPA (**E**) Bar graphs showing the average Ca^2+^ event frequency for aging mice after bath application of zero Ca^2+^ aCSF (left graph) or after 15 min incubation with 20 µM CPA (right graph) (**F**) Bar graph of average % decrease in Ca^2+^ event frequency in aging mice exposed to either zero Ca^2+^ aCSF or CPA. For zero Ca^2+^ in young mice, n = 64 ROIs and 20 blood vessels from 5 mice. For zero Ca^2+^ in aging mice, n = 57 ROIs and 26 blood vessels from 7 mice. For CPA experiments in young mice, n = 21 ROIs and 11 blood vessels from 5 mice. For CPA experiments in aging mice, n = 24 ROIs and 11 blood vessels from 6 mice Error bars are S.E.M and *p* values are based on Wilcoxon signed ranked tests.

**Figure 5.**
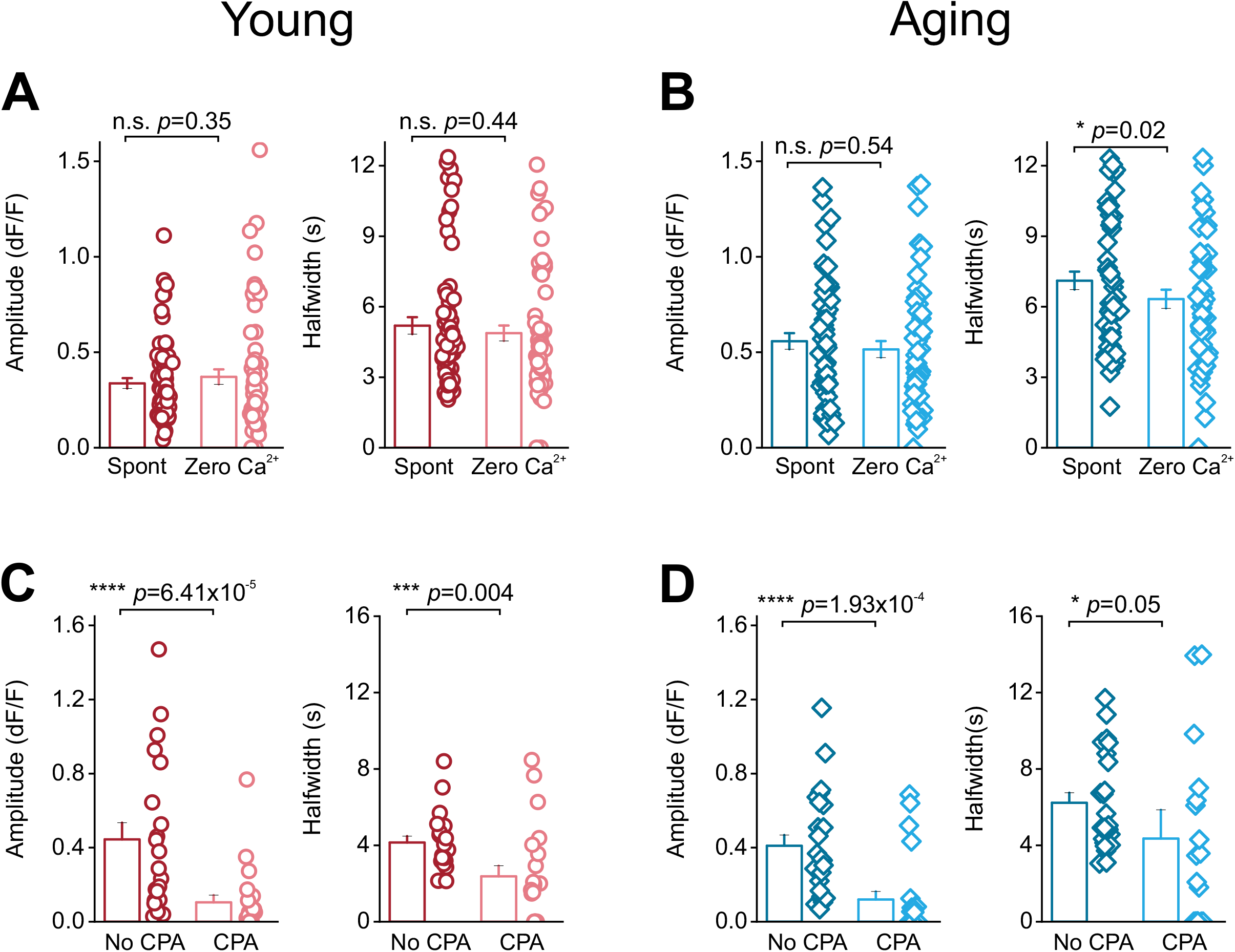
CPA alters endfoot Ca^2+^ event amplitude and duration in young and aging mice. (**A**) Bar graphs showing amplitude (left) and half width (right) for Ca^2+^ events with zero Ca^2+^ in young mice (**B**) Bar graphs showing amplitude (left) and half width (right) for Ca^2+^ events with zero Ca^2+^ in aging mice (**C**) Bar graphs showing amplitude (left) and half width (right) for Ca^2+^ events with CPA in young mice (**D**) Bar graphs showing amplitude (left) and half width (right) for Ca^2+^ events with CPA in aging mice. For zero Ca^2+^ experiments, young mice, n = 64 ROIs and 20 blood vessels from 5 mice. For aging mice, n = 57 ROIs and 26 blood vessels from 7 mice. For CPA experiments, young mice, n = 21 ROIs and 11 blood vessels from 5 mice and aging mice, n = 24 ROIs and 11 blood vessels from 6 mice. Error bars are S.E.M and *p* values are based on Wilcoxon signed ranked tests.

For aging mice, zero Ca^2+^ aCSF did not affect Ca^2+^ event frequency (Figure 4D and E; supplemental movie 2) or amplitude but caused a small decrease in half width (Figure 5B). However, following CPA treatment, DLS brain slices of aging mice showed a dramatic 75% reduction in Ca^2+^ event frequency (Figure 4D and E). Aging mice also demonstrated a 71% decrease in amplitude and a 30% decrease in half width of the Ca^2+^ events that persisted after CPA application (Figure 5D).

These data show that while young mice rely on both extracellular and intracellular Ca^2+^ stores, aging mice almost exclusively rely on ER Ca^2+^ for generating spontaneous endfoot Ca^2+^ events in the DLS (Figure 4C and F).

### Astrocytic endfeet in the DLS of aging mice demonstrate significant reductions in CALR

Based on the finding that aging mice exclusively rely on ER Ca^2+^ for generating endfoot Ca^2+^ events (Figures 4 and 5), we rationalized that aging may be associated with an alteration in ER Ca^2+^ buffering. To test this hypothesis, young and aging striatal brain sections were co-immunostained for the major ER-localized Ca^2+^ buffering protein, CALR (Nakamura et al., 2001), and the astrocytic endfoot marker, AQP4. In both young and aging mice, AQP4 labeling clearly outlined BVs, while CALR staining appeared as punctate structures throughout AQP4 stained endfeet as well as the surrounding neuropil (Figure 6A). CALR positive punctae that co-localized with AQP4 were used to generate ROIs in BVs for analysis. We found that the number of CALR punctae in AQP4 co-stained endfeet decreased by ∼42% from 250.23 ± 17.6 per BV in young mice to 147.28 ± 8.23 per BV in aging mice (Figure 6B). In addition, when compared to young mice, CALR intensity in endfeet was reduced by ∼20% (Figure 6C). This reduction in CALR intensity in aging mice was not restricted to endfeet since we observed a general 15% decrease in CALR intensity throughout the neuropil of the DLS in aging mice (Figure 6D). Thus, when compared to young mice, aging mice demonstrate a significant reduction in expression of the major Ca^2+^ buffer, CALR within the neuropil and astrocytic endfeet of the DLS.

**Figure 6.**
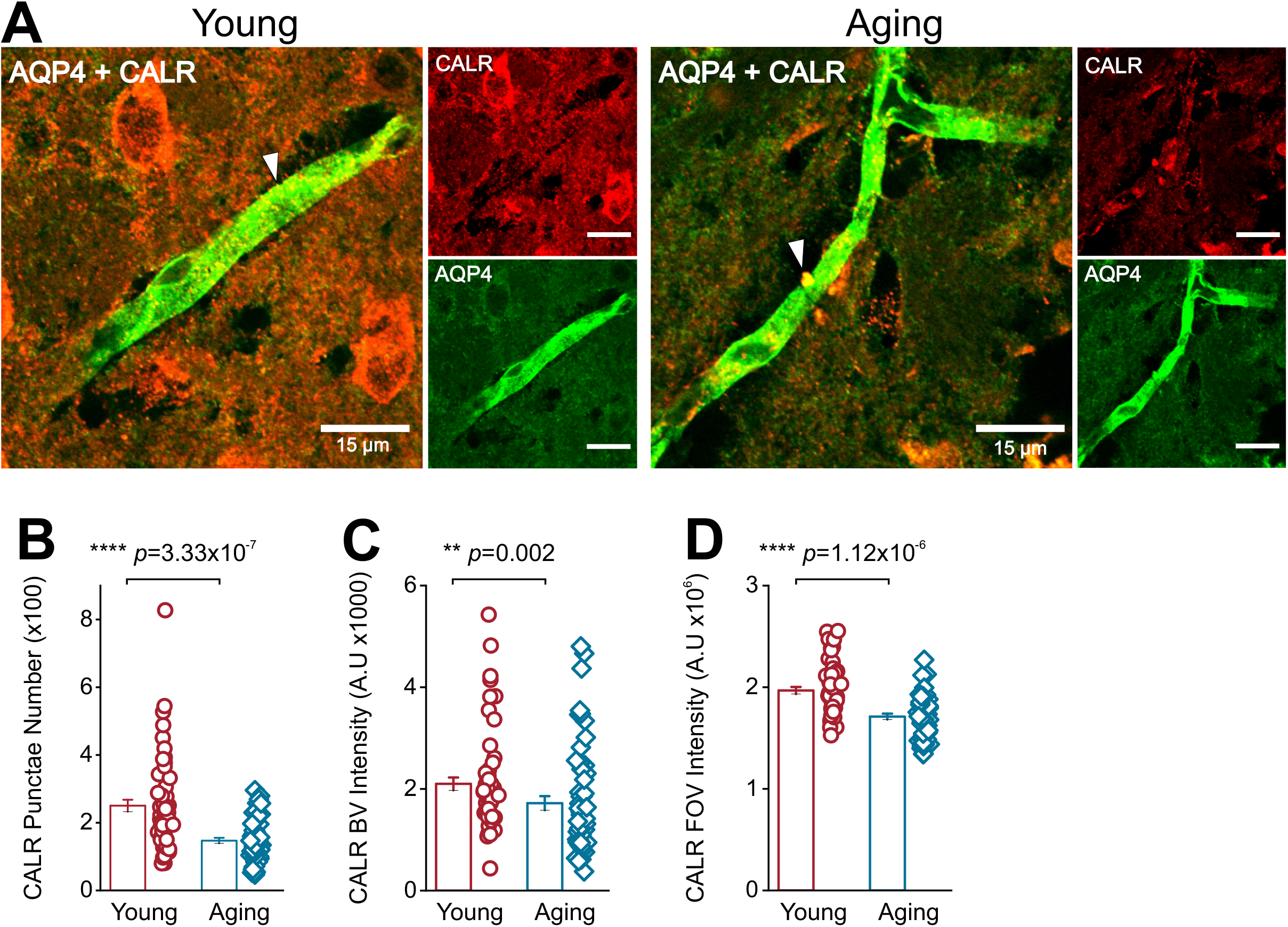
Aging reduces CALR expression levels in astrocyte endfeet and in the neuropil of the DLS. (**A**) Z-stack of representative confocal images from the DLS of young (left) and aged (right) mice co-labeled with AQP4 (green) and CALR (red), scale bar = 15 μm. The white arrows point to individual CALR punctae that colocalize with AQP4 staining. (**B**) Bar graph showing the number of CALR punctae in the DLS astrocytic endfeet of young and aging mice (C) Bar graph showing CALR intensity within AQP4 labeled BVs of young and aging mice (D) Bar graph comparing CALR intensity in the entire neuropil of the DLS from young and aging mouse brain sections. For young and aging mice n = 18 blood vessels from 3 striatal sections per mouse and 3 mice per group. Error bars are S.E.M, *p* values are based on Mann-Whitney tests.

### Aging increases the frequency of spontaneous Ca^2+^ influx into endfoot mitochondria

Since the ER serves as a major source for Ca^2+^ influx into astrocytic mitochondria in the DLS (Huntington and Srinivasan, 2021), we rationalized that an aging-related reduction of CALR within astrocytic endfeet would likely alter Ca^2+^ influx into endfoot mitochondria.

To specifically measure Ca^2+^ influx into endfoot mitochondria in the DLS, young and aging mice were stereotaxically injected with a previously described construct, AAV2/5-GfaABC1D-mito7-GCaMP6f (Huntington and Srinivasan, 2021), and confocal imaging was performed in live striatal brain slices from young and aging mice with TL as a marker for BVs (Figure 7A and B). Endfoot mitochondria in young and aging mice displayed robust spontaneous Ca^2+^ influx events (Figure 7B and C; supplementary movies 5 and 6). However, when compared to young mice, aging mice showed a 35% decrease in the number of endfoot mitochondrial Ca^2+^ event ROIs and a ∼50% increase in the size of ROIs (Figure 7D).

**Figure 7.**
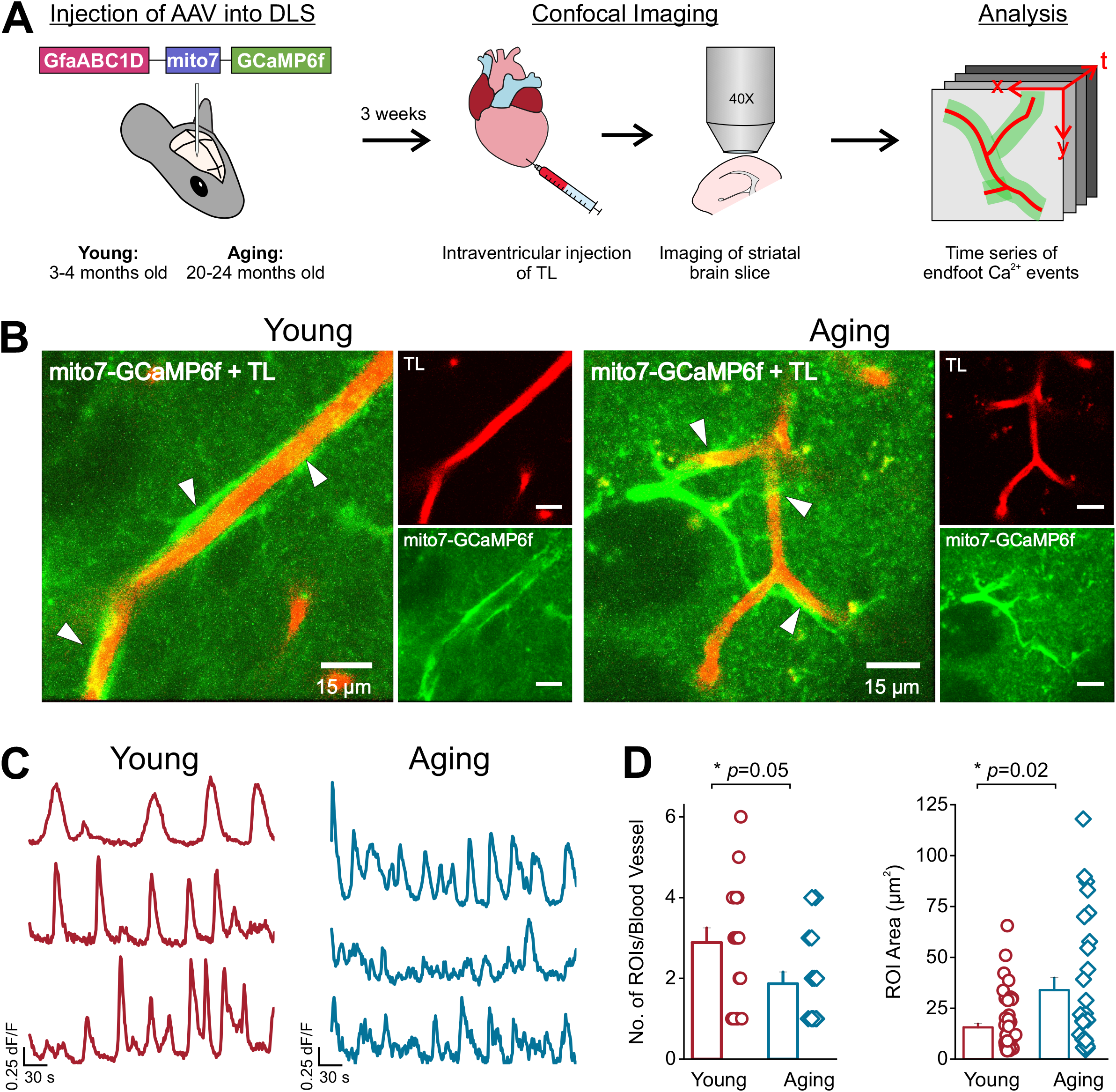
Astrocyte endfoot expression of mito7-GCaMP6f in young and aging mice. (**A**) AAV-GfaABC1D-mito7-GCaMP6f was injected into the DLS of young and aging mice. 3 weeks later, intraventricular injection of tomato lectin (TL) was performed, and striatal brain sections were collected for recording and measuring Ca^2+^ influx events in endfoot mitochondria within the DLS. (**B**) Representative t-stacks of mitochondrial Ca^2+^ influx events detected by mito7-GCaMP6f (green) in young and aging astrocyte endfeet immediately adjacent to TL labeled blood vessels (red) in the DLS, scale bar = 15 μm. White arrows point to areas where mitochondrial endfoot Ca^2+^ events initiated. (**C**) Representative endfoot mitochondrial Ca^2+^ influx event traces from young (red) and aging (blue) mice. (**D**) Bar graphs showing the number (left) and area (right) of endfoot mitochondrial Ca^2+^ event ROIs in young and aging mice. For young mice, n = 52 ROIs and 18 blood vessels from 3 mice. For aging mice, n = 28 ROIs and 16 blood vessels from 5 mice. Error bars are S.E.M and all *p* values are based on Mann-Whitney tests.

There were also significant aging-related changes in the kinetics of mitochondrial Ca^2+^ influx events. When compared to young mice, aging mice demonstrated a dramatic 50% increase in the frequency of endfoot mitochondrial Ca^2+^ influx (Figure 8A). Cumulative frequency distributions of DLS endfoot Ca^2+^ events showed that when compared to young mice with an equal distribution of mitochondrial Ca^2+^ influx across the entire frequency range (0.2 to 2.0 events/min), the vast majority (∼93%) of Ca^2+^ events in aging mice occurred at a significantly higher range (0.2 to 2.4 events/min) (Figure 8B). Despite these changes, the interval between independent endfoot mitochondrial Ca^2+^ events across BVs from multiple mice were similar in young and aging mice (Figure 8C). For both young and aging mice, there were no significant alterations in the amplitude or half width of spontaneous mitochondrial Ca^2+^ influx events (Figure 8D). Thus, aging-related increases in ER Ca^2+^ specifically caused a dramatic increase in the frequency of Ca^2+^ influx in endfoot mitochondria of the DLS.

**Figure 8.**
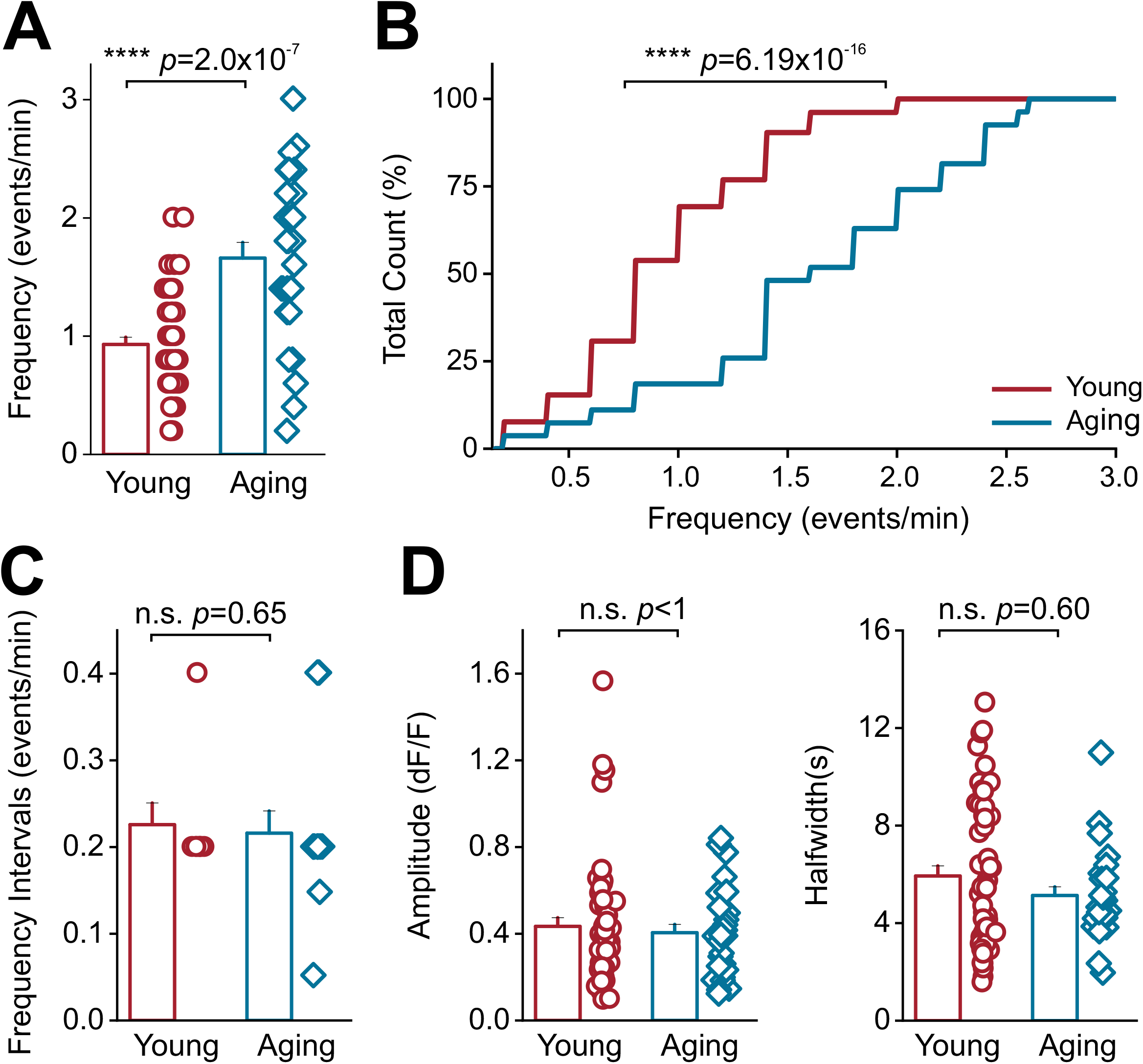
Aging increases the frequency of endfoot mitochondrial Ca^2+^ events. (**A**) Bar graph of mitochondrial Ca^2+^ event frequencies in the endfeet of young and aging mice (**B**) Cumulative distribution of mitochondrial Ca^2+^ event frequencies in young and aging mice (**C**) Plot of mitochondrial frequency intervals in young and aging mice (**D**) Amplitude (left) and half width (right) of mitochondrial Ca^2+^ events in young and aging mice. For young mice, n = 64 ROIs and 20 blood vessels from 5 mice. For young mice, n = 52 ROIs and 18 blood vessels from 3 mice. For aging mice, n = 28 ROIs and 16 blood vessels from 5 mice. Error bars are S.E.M and *p* values in panels A, C, and D are based on Mann-Whitney tests. For the cumulative frequency plot in panel B, the *p* value is based on a Kolmogorov-Smirnov test.

### Ca^2+^ influx into endfoot mitochondria of the DLS in aging mice requires ER Ca^2+^ stores

The observed increase in ER Ca^2+^ stores within astrocytic endfeet of aging mice, along with a significant aging-related increase in the frequency of Ca^2+^ influx into endfoot mitochondria led us to ask if ER Ca^2+^ stores are a primary source for Ca^2+^ influx into astrocytic endfoot mitochondria for both young and aging mice. To test this idea, we sequentially depleted extracellular, followed by intracellular Ca^2+^ sources in DLS slices. In each case, we measured the kinetics of Ca^2+^ influx into endfoot mitochondria within the DLS of young and aging mice. Extracellular Ca^2+^ depletion with zero Ca^2+^ aCSF did not alter the amplitude of endfoot mitochondrial Ca^2+^ influx in young and aging mice (Figure 9B and D; supplementary movies 5 and 6). Young mice demonstrated no change in Ca^2+^ event frequency and a small decrease in the half width of mitochondrial Ca^2+^ signals (Figure 9B). For aging mice, we observed a small decrease in the frequency, but no change in the half width of mitochondrial Ca^2+^ signals (Figure 9D). In contrast, depletion of ER Ca^2+^ stores with CPA resulted in a dramatic and significant ∼80% decrease in the frequency, amplitude and half width of mitochondrial endfoot Ca^2+^ influx in young mice (Figure 10A and B; supplementary movies 7 and 8). In aging mice, we observed a significant 77% decrease in frequency, 65% decrease in amplitude, and 59% decrease in event half width (Figure 10C and D). These data confirm that the ER is indeed a primary source for Ca^2+^ influx into astrocytic endfoot mitochondria for both young and aging mice.

**Figure 9.**
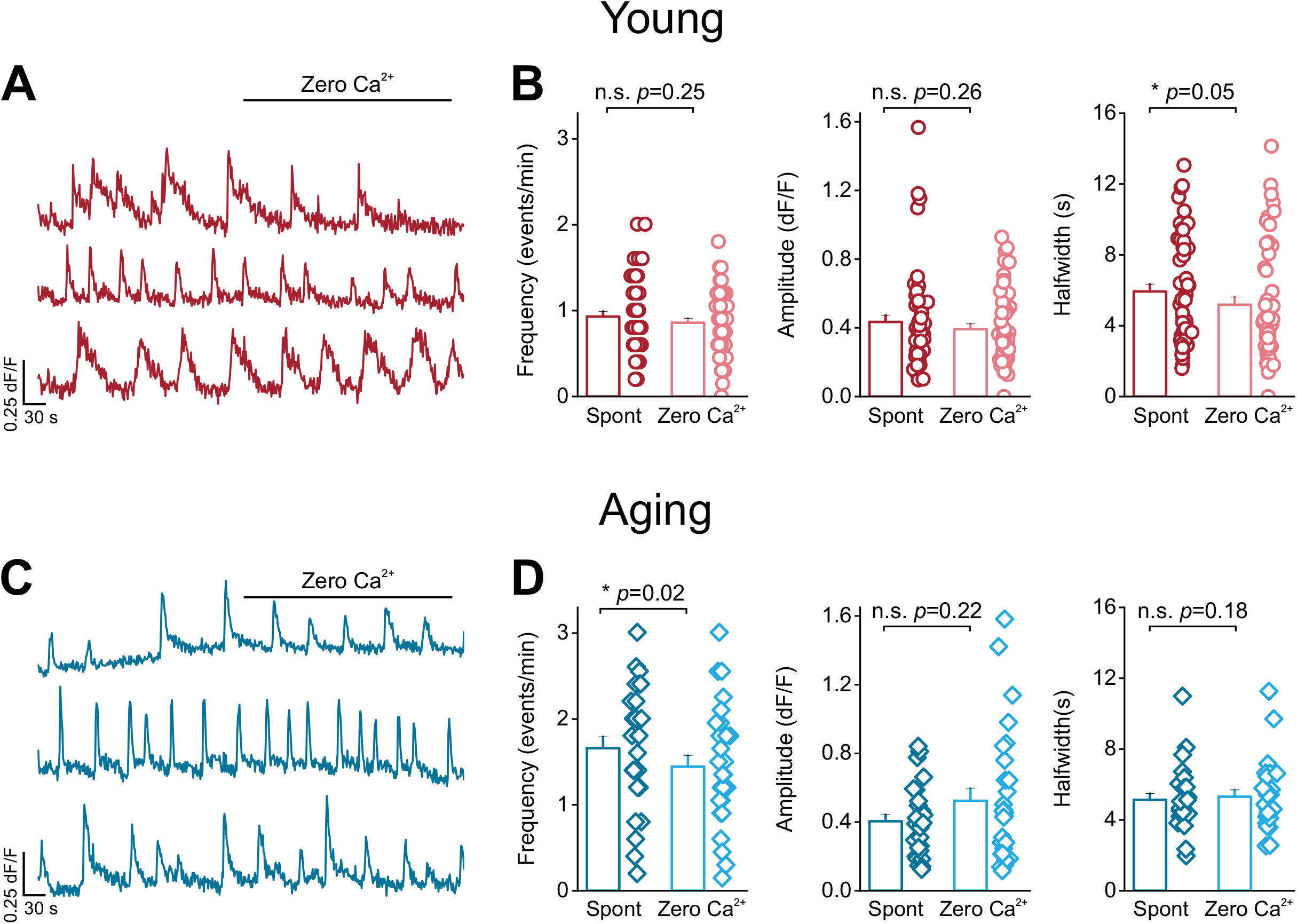
Endfoot mitochondria in young and aging mice do not rely on extracellular Ca^2+^. (**A**) Representative mitochondrial Ca^2+^ event traces from young mice before and after bath application of zero Ca^2+^ aCSF (**B**) Bar graphs showing frequency (left), amplitude (middle), and half width (right) for mitochondrial Ca^2+^ events recorded in young mice with and without zero Ca^2+^ aCSF (**C**) Representative mitochondrial Ca^2+^ event traces from aging mice before and after bath application of zero Ca^2+^ aCSF (**D**) Bar graphs showing frequency (left), amplitude (middle), and halfwidth (right) for mitochondrial Ca^2+^ events recorded in aging mice with and without zero Ca^2+^ aCSF. For young mice, n = 52 ROIs and 18 blood vessels from 3 mice. For aging mice, n = 28 ROIs and 16 blood vessels from 5 mice. Error bars are S.E.M and all *p* values are based on Wilcoxon signed ranked tests.

**Figure 10.**
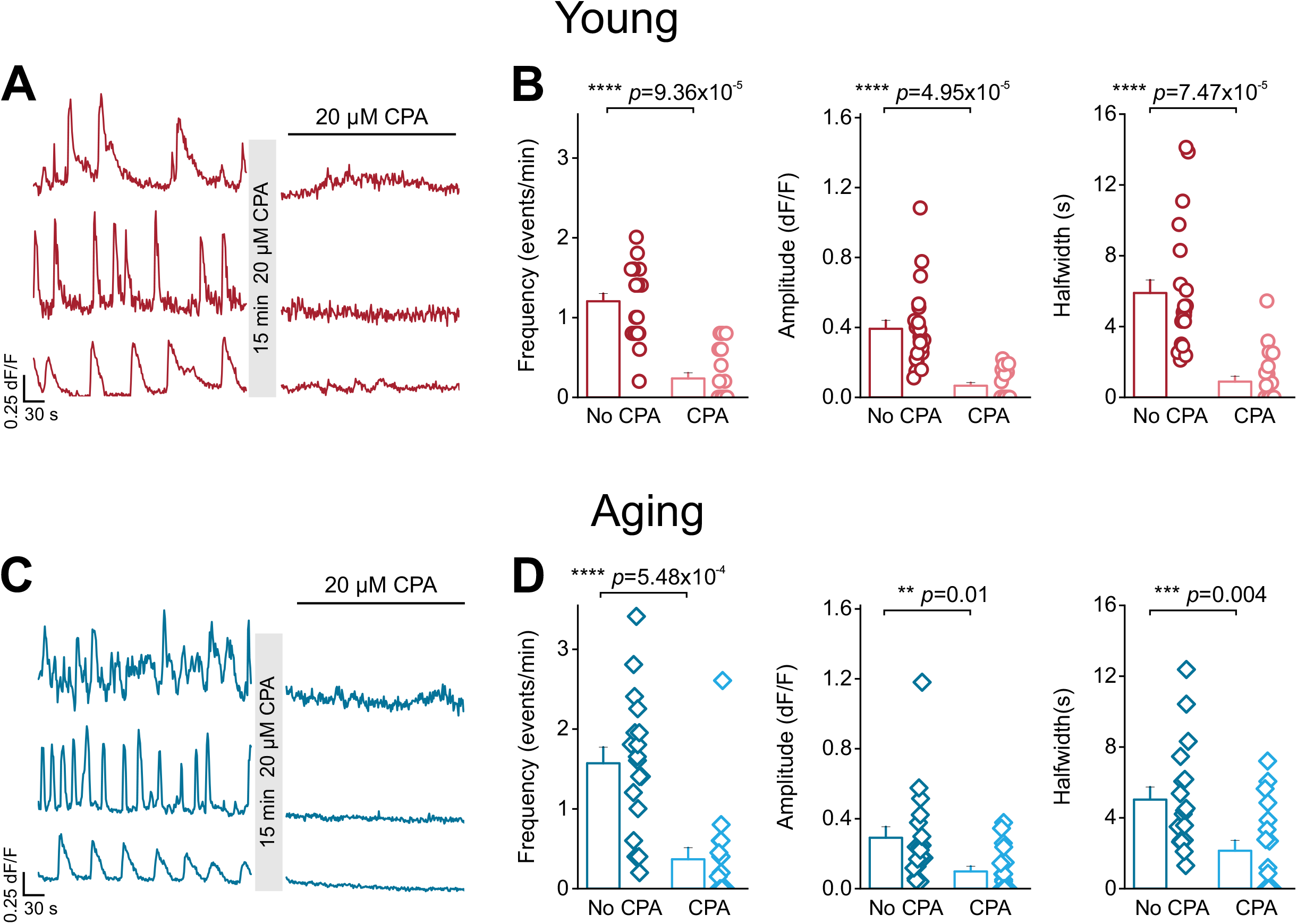
Endfoot mitochondria in young and aging mice depend on intracellular Ca^2+^ stores. (**A**) Representative mitochondrial Ca^2+^ event traces from young mice before and after bath application of CPA (**B**) Bar graphs showing frequency (left), amplitude (middle), and half width (right) for mitochondrial Ca^2+^ events recorded in young mice with and without CPA (**C**) Representative mitochondrial Ca^2+^ event traces from aging mice before and after bath application of CPA (**D**) Bar graphs showing frequency (left), amplitude (middle), and halfwidth (right) for mitochondrial Ca^2+^ events recorded in aging mice with and without CPA. For young mice, n = 22 ROIs and 9 blood vessels from 3 mice. For aging mice, n = 18 ROIs and 11 blood vessels from 5 mice All errors are SEM; *p* values are based on Wilcoxon signed ranked tests.

## Discussion

Aging is associated with subtle changes in neuronal and astrocytic Ca^2+^ homeostasis (Guerra-Gomes et al., 2018; Nikoletopoulou and Tavernarakis, 2012; Toescu and Verkhratsky, 2007). In addition, neurovascular impairment is known to initiate neurodegeneration (Cabezas et al., 2014; Cai et al., 2017; Govindpani et al., 2019; Grammas et al., 1999; Ott et al., 2018; Sweeney et al., 2019; Yu et al., 2020; Zhou et al., 2020), and aging alters astrocytic gene expression (Boisvert et al., 2018). These seemingly unrelated findings point to the idea that aging could alter Ca^2+^-mediated coupling between astrocytic endfeet and the neurovasculature, eventually leading to neurodegeneration. In this context, we asked if astrocytic endfeet in the DLS of aging mice display changes in expression of the Ca^2+^ buffering protein, CALR, and alterations in the sources and kinetics of astrocytic endfoot Ca^2+^ signals.

We found that aging astrocytic endfeet in the DLS demonstrate a significant reduction in CALR (Figure 6), which is the primary endogenous buffer for ER-localized Ca^2+^. This reduction in CALR is associated with alterations in multiple aspects of spontaneous cytosolic and mitochondrial Ca^2+^ events within astrocytic endfeet (Figures 1,2,3,7 and 8). Based on these data, we infer that the aging-related reduction of CALR expression is upstream of alterations in endfoot Ca^2+^ signals. Multiple lines of evidence support our interpretation of the data: (**i**) Astrocytic endfeet contain ER and mitochondria (Boulay et al., 2017; Gobel et al., 2020), (**ii**) The generation of spontaneous endfoot Ca^2+^ signals in aging mice exclusively depends on ER rather than extracellular Ca^2+^ stores (Figure 4C and F), and (**iii**) As previously shown by Biwer *et al*, conditional deletion of CALR from endothelial cells in aging mice increases the spatial spread of spontaneous endothelial Ca^2+^ signals and impairs Ca^2+^ mobilization (Biwer et al., 2020).

Similar to the Biwer *et al* study which was conducted in mice with a conditional knockout of CALR in endothelial cells (Biwer et al., 2020), the current study shows that reduced CALR expression in the astrocytic endfeet of aging mice is associated with an increase in the velocity of expansion of DLS endfoot Ca^2+^ signals (Figure 3). Since CALR is known to regulate Ca^2+^ mobilization within the ER (Nakamura et al., 2001), our findings suggest that the observed changes in Ca^2+^ signal kinetics likely stem from the aging-related reduction of CALR within astrocytic endfeet. In accordance with this rationale, we observed very specific aging-related changes in the kinetics of cytosolic astrocytic endfoot Ca^2+^ signals. There was an increase in the number of low frequency endfoot Ca^2+^ signals in aging mice (Figure 2B) accompanied by a nearly 2-fold increase in the amplitude as well as an increase in the duration of Ca^2+^ signals (Figure 2D). These data suggest that aging mice possess a generalized dysregulation of Ca^2+^ signaling in astrocytic endfeet of the DLS.

Our finding that endfoot Ca^2+^ signals in aging mice exclusively depend on ER stores (Figure 4F) strongly suggests that the observed changes in cytosolic Ca^2+^ signal kinetics likely reflect Ca^2+^ efflux from the ER into the cytosol of astrocytic endfeet. The ER in astrocytic endfeet is morphologically and functionally coupled with mitochondria (Boulay et al., 2017; Gobel et al., 2020), enabling mitochondria to sequester Ca^2+^ released from the ER. Therefore, one downstream target of aging-related alterations in Ca^2+^ efflux from the ER of astrocytic endfeet would be changes in mitochondrial Ca^2+^ signals. In line with this rationale, we showed that when compared to young mice, aging-related alterations in endfoot Ca^2+^ efflux from the ER of aging mice was associated with a concomitant increase in the frequency of Ca^2+^ influx events into endfoot mitochondria (Figure 8A and B). The finding that mitochondrial Ca^2+^ influx increases in frequency with a corresponding decrease in the frequency of cytosolic Ca^2+^ signals suggests a general aging-related dysregulation of functional coupling between the ER and mitochondria in astrocytic endfeet of the DLS. Given the importance of normal AQP4 localization and function in maintaining BBB integrity (Vella et al., 2015), one downstream consequence of reduced CALR expression and altered Ca^2+^ signals in aging astrocytic endfeet may be the gradual loss of AQP4, resulting in a loss of BBB integrity and a consequent increase in the risk for neurodegeneration. In support of this idea, a reduction in CALR expression has been linked to the loss of motor neurons during amyotrophic lateral sclerosis (ALS) (Bernard-Marissal et al., 2012), and physical and functional changes in the NVU have been established as an important contributor to neurodegenerative disorders such as AD and PD (Cabezas et al., 2014; Cai et al., 2017; Govindpani et al., 2019; Grammas et al., 1999; Ott et al., 2018; Sweeney et al., 2019; Yu et al., 2020; Zhou et al., 2020). Apart from altering BBB integrity, subtle changes in the localization of Ca^2+^ fluxes within the aging astrocytic endfoot could have important consequences for vascular endothelial cell function. Conceivably, altered Ca^2+^ signals could change the activation of large-conductance Ca^2+^ activated K^+^ (BK) channels in aging endfeet (Masamoto et al., 2015), thereby disrupting K^+^-dependent endothelial cell-mediated capillary to arteriole ascending dilation within the brain (Bagher and Segal, 2011; Longden et al., 2014; Longden et al., 2017).

In summary, we show that an aging-related reduction in CALR expression within astrocytic endfeet results in the exclusive dependence of endfoot Ca^2+^ signals on ER Ca^2+^, which results in abnormal Ca^2+^ coupling between the ER and mitochondria. Based on our data, we propose that altered Ca^2+^ buffering and aging-related dysregulation in endfoot Ca^2+^ signaling could conceivably alter the localization of AQP4 within astrocytic endfeet, thereby compromising BBB integrity, as well as NVU coupling. Taken together, our findings have important implications for understanding how BBB and NVU dysfunction in the aging brain can trigger, and possibly sustain neurodegenerative processes.

## Acknowledgements

This work was partially funded by R01HL155618 to PB and an American Diabetes Association Junior Faculty Development Award, 1-19-JDF-111 to PB and by a National Institutes of Health (NIH) research grant, R01NS115809 to RS.

## Abbreviations

BBB: blood brain barrier
NVU: neurovascular unit
BV: blood vessel
CALR: calreticulin
ER: endoplasmic reticulum
AAV: adeno-associated virus
PD: Parkinson’s disease
AD: Alzheimer’s disease
aCSF: artificial cerebrospinal fluid
DLS: dorsolateral striatum
AQP4: aquaporin 4

## Notes

**Conflict of interest statement:** Authors declare no competing financial interests.

### Competing Interest Statement

The authors have declared no competing interest.

